# Training neural networks to recognize speech increased their correspondence to the human auditory pathway but did not yield a shared hierarchy of acoustic features

**DOI:** 10.1101/2021.01.26.428323

**Authors:** Jessica A.F. Thompson, Yoshua Bengio, Elia Formisano, Marc Schönwiesner

## Abstract

The correspondence between the activity of artificial neurons in convolutional neural networks (CNNs) trained to recognize objects in images and neural activity collected throughout the primate visual system has been well documented. Shallower layers of CNNs are typically more similar to early visual areas and deeper layers tend to be more similar to later visual areas, providing evidence for a shared representational hierarchy. This phenomenon has not been thoroughly studied in the auditory domain. Here, we compared the representations of CNNs trained to recognize speech (triphone recognition) to 7-Tesla fMRI activity collected throughout the human auditory pathway, including subcortical and cortical regions, while participants listened to speech. We found no evidence for a shared representational hierarchy of acoustic speech features. Instead, all auditory regions of interest were most similar to a single layer of the CNNs: the first fully-connected layer. This layer sits at the boundary between the relatively task-general intermediate layers and the highly task-specific final layers. This suggests that alternative architectural designs and/or training objectives may be needed to achieve fine-grained layer-wise correspondence with the human auditory pathway.

**Highlights:**

- Trained CNNs more similar to auditory fMRI activity than untrained
- No evidence of a shared representational hierarchy for acoustic features
- All ROIs were most similar to the first fully-connected layer
- CNN performance on speech recognition task positively associated with fmri similarity

## 1. Introduction

The use of deep neural networks (DNNs) as models of biological neural networks has been discussed as an opportunity for synergy between neuroscience and artificial intelligence (Barrett et al., 2019, Marblestone et al., 2016, Richards et al., 2019). The paradigm of comparing DNN activity to neural activity has been most thoroughly explored in research on the primate visual system. Seminal work by DiCarlo & Cox proposed that visual object recognition is accomplished via successive layers of nonlinear transformations that effectively *untangle* visual inputs, linearizing the boundaries between object manifolds (DiCarlo and Cox, 2007). Similar language has been used to describe how DNNs accomplish recognition tasks (Bengio et al., 2013). Several studies have now reported that state-of-the-art (SOTA) machine learning systems, trained only to maximize their performance on a specific task, without any explicit goal to mimic neural activity, appear to learn representations that are similar to those found in the brains of animals engaged in a similar task (Kriegeskorte, 2015). For example, the output layer of Alexnet (Krizhevsky and Hinton, 2012) has been found to be highly predictive of spiking responses to natural images in inferior temporal cortex and intermediate layers to be highly predictive of V4 responses (Cadieu et al., 2014, Yamins et al., 2014). Similar comparisons have been made between modern convnets and the human visual system as recorded with functional magnetic resonance imaging (fMRI) (Khaligh-Razavi and Kriegeskorte, 2014, Agrawal et al., 2014, Eickenberg et al., 2017, Guclu and van Gerven, 2016). The most convincing demonstration that modern convnets learn representations that are meaningful to neurons in the primate visual system is work from Bashivan et al. (2019) showing that task-optimized DNNs can be used to control the activity of macaque V4 neurons. They found that stimuli synthesized to maximally activate specific units in the DNN also drove activity of matched sites in V4 well beyond their maximum firing rate in response to natural images.

Comparisons of DNNs to biological sensory pathways often come with claims of shared representation hierarchy. Regions of interest (ROIs) along some pathway are mapped to layers of a DNN based on their similarity. Early layers in the network tend to be more similar to early ROIs in the pathway and late layers to late ROIs (Cichy et al., 2016, Güçlü and van Gerven, 2015). These results suggest that DNNs are not just learning representations that are similar to single regions, but rather that they constitute models of an entire hierarchy of sensory processing. However, not all studies have found evidence of shared hierarchy. Cadena et al. (2019) compared representations at several layers of a convnet trained on ImageNet to neural activation in the mouse visual cortex. While they found their network outperformed classical predictive models, they found no evidence for a shared hierarchy and no benefit over a random network whose weights had never been trained. The authors suggest that networks trained on more ethologically valid tasks may be required to capture the functional organization of the rodent visual cortex.

Relatively few experiments have compared DNNs trained on acoustic tasks to biological auditory systems. Kell et al. (2018) trained convnets on speech and music tasks and compared their learned representations to fMRI responses in human auditory cortex. They found that intermediate DNN representations explained more variance in auditory cortex responses than a spectrotemporal modulation-based baseline model. To assess the existence of a shared hierarchy, they looked only at voxels that showed a reliable response to sound and layers of their network which were predictive of voxel activity across auditory cortex. They found that the most predictive layers of primary auditory cortex were intermediate layers, while the most predictive layers of secondary auditory cortex were deeper layers. From this, they conclude that the hierarchical distinction between primary and secondary auditory cortex is mirrored in their convnet (Kell et al., 2018). Güçlü et al. also reported evidence for a shared hierarchy in human auditory cortex, but they only analyzed the superior temporal gyrus (STG). They used representational similarity analysis (RSA) to compare representations learned in a DNN trained to predict tags from excerpts of musical audio.1 They found a gradient of complexity across STG where anterior voxel clusters were more similar to early layers while posterior voxel clusters were more similar to late layers (Güçlü et al., 2016). While both of the above studies report evidence for a shared hierarchy between human auditory cortex and DNNs trained on sound, they report different spatial patterns of similarity gradients.

Several different analysis tools are used to compare representations. The ultimate goal of these analyses is to quantify the similarity of two representations, but similarity is an ambiguous term that must be defined by the experimenter. In many of the aforementioned studies, an encoding analysis is performed where firing rate or voxel activity is predicted by a regularized linear model of the neural network activity. According to this approach, a representation is similar to another to the extent that it can be linearly predicted from the other. There are other notions of representational similarity that have been explored to study DNNs. Singular value canonical correlation analysis (SVCCA) and projection-weighted canonical correlation analysis (pwCCA) have been used to characterize how network representations change over training, to compare representations in different architectures, and to understand the difference between networks that memorize and networks that generalize (Raghu et al., 2017, Morcos et al., 2018). Kornblith et al. recently proposed that, given two networks of identical architecture and training, differing only in their random initialization, a meaningful notion of similarity should find their corresponding layers to be most similar (i.e. layer 1 in network A should be most similar to layer 1 in network B). Of the tested metrics, which included SVCCA, pwCCA and linear regression, Centered Kernel Alignment (CKA) was the only method which found that corresponding layers were most similar to each other, achieving an accuracy of 99.3% on the layer identification task. The next best metric, linear regression, achieved only 45.4%. This result may be related to the fact that CKA is only invariant to orthogonal transformations and isotropic scaling, unlike canonical correlation analysis (CCA), which is invariant to any linear invertible transformation, and linear regression, which is invariant to any linear invertible transformation of the predicted variables (Kornblith et al., 2019). Representational similarity analysis (RSA) (Kriegeskorte et al., 2008), commonly employed in fMRI analysis, is similar to CKA with a linear kernel except that CKA is based on dot-product similarity and RSA typically uses correlation-based metrics. CKA provides a general framework with interpretable units, proven convergence rates, and the option to use different kernels.

Here, we use CKA to quantify the similarity between representations learned in convnets trained on speech and activity throughout the human auditory pathway during speech listening, as measured with 7-Tesla (7T) fMRI. The high spatial resolution of 7T fMRI allows us to simultaneously measure activity from auditory cortex as well as subcortical auditory regions, which are often omitted from auditory fMRI analyses due to their small size. Since significant auditory processing occurs in brainstem and midbrain regions, this provides us with several distinct regions with a relatively known connectivity structure with which to compare the convnet representations. To the best of our knowledge, ours is the first study to compare DNN representations to activity throughout the human subcortical and cortical auditory pathway. If there exists a shared hierarchy between the convnets and the human auditory pathway, the pattern of similarity should at least distinguish between cortical and subcortical regions. We visualized the results of the similarity analysis as similarity matrices with network layers as the rows and auditory ROIs as the columns. Evidence of a shared hierarchy would manifest as a diagonal pattern in one such similarity matrix, where shallower layers are more similar to early regions and deeper layers more similar to later regions. While we found that our trained networks were more similar to the brain than an untrained network, we found no such diagonal pattern. Instead we found that, on average, nearly all ROIs are most similar to the first fully-connected layer.

## 2. Material and methods

### 2.1. Participants

Six healthy participants (aged 28–31, three women, three men) with normal hearing and no known neurological disorders were recruited to participate. All participants provided written informed consent prior to the first MRI session. All participants also consented to their data being made publicly available.2 The native languages of the participants were English (one subject), German (three participants) and Dutch (two participants).

### 2.2. Experimental Stimuli

To facilitate comparison with the convnets, we selected utterances from the same corpus that the networks were trained on. Such a comparison is complicated by the fact that, although the networks were only trained on phonetic labels, human listeners will perceive the meaning and higher-level structure of speech, even if not instructed to do so. Therefore, to make the experimental conditions as similar as possible for both human and network listeners, we transformed the natural speech to remove higher-level structure while preserving the original phonemes. This quilting procedure, described below, allowed us to focus our comparison on representational transformations only up to the sub-word level in both the convnets and the human auditory system.

The audio corpora from which the stimuli were constructed were the same datasets that were used in (Thompson et al., 2019a) and (Thompson et al., 2019b), which are owned by Nuance Communications. Each of the three datasets, one for English, Dutch and German, contained 64-83 hours of spoken text read by several native speakers in a quiet room. The datasets also included phonetic transcriptions established in a forced alignment with text transcriptions.

The quilting procedure, adapted from (Overath et al., 2015), chops a sound file into small segments and reorders the segments according to a heuristic designed to hide the *seams* of the quilt (the segment boundaries).3 A random segment is chosen as the first segment in the quilt. Subsequent segments are chosen to best match the segment-to-segment boundaries in the chochleogram of the original audio. In this way, temporal patterns longer than the segment length are destroyed while minimizing the artefacts introduced by reordering the segments.

Instead of using fixed segment lengths, as in (Overath et al., 2015), we used the provided phonetic boundaries to divide the speech into variable length segments containing single phonemes. The resulting quilts are out-of-order sequences of phonemes, preserving phonetic information while destroying the words and semantic content of the speech. The larger the input corpus relative to the desired quilt length, the more effectively the seams of the quilt will be hidden. Therefore, we selected the 60 speakers (30 women and 30 men) with the longest set of utterances in each language. Given all the utterances from a single speaker as input, the quilting procedure generated a one-minute quilt. The experimental stimuli consisted of 180 one-minute speech quilts (60 per English, Dutch and German). The final stimuli were filtered to account for the frequency response profile of the foam-tip earphones over which the stimuli were presented in the scanner.

### 2.3. Experimental Protocol

The experimental procedures were approved by the ethics committee of the Faculty for Psychology and Neuroscience at Maastricht University (approval code ERCPN-167_09_05_2016). Magnetic resonance images were collected over two sessions on separate days, each consisting of 10 functional runs. Nine speech quilts were presented in each run, grouped into blocks of three quilts from the same language. Within a block, the quilts were presented one after another with no interruption. Blocks were separated by short periods of rest which were sometimes followed by a question asking participants to identify the language of the speech presented in the preceding block. The purpose of this question was to ensure that participants were awake and paying attention to the stimuli. Participants used a button box to indicate their response. To save time, this vigilance question was not asked after every block. However, the design was such that the participants could not easily predict whether they would be questioned and so had to pay attention during every block. Each run contained one block for each language. The stimuli were presented in a different pseudo-random order for each participant.

### 2.4. MRI Acquisition Parameters

Images were acquired at Maastricht University, Maastricht, Netherlands on a 7T Siemens MAGNETOM scanner (Siemens Medical Solutions, Erlangen, Germany), with 70 mT/m gradients and a head RF coil (Nova Medical, Wilmington, MA, USA; single transmit, 32 receive channels). Foam pads were used to minimize head motion.

#### 2.4.1. Anatomical

At the start of each session, a T1-weighted (T1w) image and a proton density weighted (PDw) image were acquired using a 3D MPRAGE sequence [voxel size=1.0mm isotropic; repetition time (TR)=2370 ms; echo time (TE)=2.31 ms; flip angle=5°; generalized auto-calibrating partially parallel acquisitions (GRAPPA)=3 (Griswold et al., 2002); field of view (FOV)=256 mm; 256 slices, phase encoding direction: anterior to posterior, inversion time (TI) for T1w only=1500 ms].

#### 2.4.2. Functional

Functional MRI data were acquired with a 2-D Multi-Band Echo Planar Imaging (2D-MBEPI) sequence (Steen Moeller et al., 2010, Setsompop et al., 2012). In order to include the entire brainstem and thalamus as well as primary and secondary auditory cortex, slices were arranged in a coronal oblique orientation (TR=1700 ms; TE=20 ms; flip angle=70°; GRAPPA=3; Multi-Band factor=2; FOV=206 mm; 1.7 mm isotropic voxels; phase encode direction inferior to superior).

### 2.5. MRI Preprocessing

The MRI preprocessing was performed using *fMRIPrep* 1.4.1 (Esteban et al. 2018a; Esteban et al. 2018b; RRID:SCR_016216), which is based on *Nipype* 1.2.0 (Gorgolewski et al. 2011; Gorgolewski et al. 2018; RRID:SCR_002502). The following description was prepared by *fMRIPrep*.

#### 2.5.1. Anatomical data preprocessing

T1-weighted (T1w) images were corrected for intensity non-uniformity (INU) with `N4BiasFieldCorrection` (Tustison et al., 2010), distributed with ANTs 2.2.0 (Avants et al., 2008, RRID:SCR004757). The T1w-reference was then skull-stripped with a *Nipype* implementation of the `antsBrainExtraction.sh` workflow (from ANTs), using OASIS30ANTs as target template. Brain tissue segmentation of cerebrospinal fluid (CSF), white-matter (WM) and graymatter (GM) was performed on the brain-extracted T1w using fast (FSL 5.0.9, RRID:SCR_002823, Zhang et al., 2001). A T1w-reference map was computed after registration of 2 T1w images (after INU-correction) using `mri_robust_template` (FreeSurfer 6.0.1, Reuter et al., 2010). Brain surfaces were reconstructed using `recon-all` (FreeSurfer 6.0.1, RRID:SCR_001847, Dale et al., 1999), and the brain mask estimated previously was refined with a custom variation of the method to reconcile ANTs-derived and FreeSurfer-derived segmentations of the cortical gray-matter of Mindboggle (RRID:SCR_002438, Klein et al., 2017). Volume-based spatial normalization to one standard space (MNI152NLin2009cAsym) was performed through nonlinear registration with `antsRegistration` (ANTs 2.2.0), using brain-extracted versions of both T1w reference and the T1w template. The following template was selected for spatial normalization: *ICBM 152 Nonlinear Asymmetrical template version 2009c* (Fonov et al. 2009, RRID:SCR_008796; TemplateFlow ID: MNI152NLin2009cAsym).

#### 2.5.2. Functional data preprocessing

For each of the 20 BOLD runs per subject (across all sessions), the following preprocessing was performed. First, a reference volume and its skull-stripped version were generated using a custom methodology of *fM-RIPrep*. The BOLD reference was then co-registered to the T1w reference using `bbregister` (FreeSurfer) which implements boundary-based registration (Greve and Fischl, 2009). Co-registration was configured with nine degrees of freedom to account for distortions remaining in the BOLD reference. Head-motion parameters with respect to the BOLD reference (transformation matrices, and six corresponding rotation and translation parameters) are estimated before any spatiotemporal filtering using `mcflirt` (FSL 5.0.9, Jenkinson et al., 2002). BOLD runs were slice-time corrected using `3dTshift` from AFNI 20160207 (Cox and Hyde, 1997, RRID:SCR_005927). The BOLD time-series, were resampled to surfaces on the following spaces: *fsaverage5*. The BOLD time-series (including slice-timing correction when applied) were resampled onto their original, native space by applying a single, composite transform to correct for head-motion and susceptibility distortions. These resampled BOLD time-series will be referred to as *preprocessed BOLD in original space*, or just *preprocessed BOLD*. The BOLD time-series were resampled into standard space, generating a *preprocessed BOLD run in [‘MNI152NLin2009cAsym’] space*. First, a reference volume and its skull-stripped version were generated using a custom methodology of *fM-RIPrep*. Several confounding time-series were calculated based on the *preprocessed BOLD*: framewise displacement (FD), DVARS and three region-wise global signals. FD and DVARS are calculated for each functional run, both using their implementations in *Nipype* (following the definitions by Power et al., 2014). The three global signals are extracted within the CSF, the WM, and the whole-brain masks. Additionally, a set of physiological regressors were extracted to allow for component-based noise correction (*CompCor*, Behzadi et al., 2007). Principal components are estimated after high-pass filtering the *preprocessed BOLD* time-series (using a discrete cosine filter with 128s cut-off) for the two *CompCor* variants: temporal (tCompCor) and anatomical (aCompCor). tCompCor components are then calculated from the top 5% variable voxels within a mask covering the subcortical regions. This subcortical mask is obtained by heavily eroding the brain mask, which ensures it does not include cortical GM regions. For aCompCor, components are calculated within the intersection of the aforementioned mask and the union of CSF and WM masks calculated in T1w space, after their projection to the native space of each functional run (using the inverse BOLD-to-T1w transformation). Components are also calculated separately within the WM and CSF masks. For each CompCor decomposition, the *k* components with the largest singular values are retained, such that the retained components’ time series are sufficient to explain 50 percent of variance across the nuisance mask (CSF, WM, combined, or temporal). The remaining components are dropped from consideration. The head-motion estimates calculated in the correction step were also placed within the corresponding confounds file. The confound time series derived from head motion estimates and global signals were expanded with the inclusion of temporal derivatives and quadratic terms for each (Satterthwaite et al., 2013). Frames that exceeded a threshold of 0.5 mm FD or 1.5 standardised DVARS were annotated as motion outliers. All resamplings can be performed with *a single interpolation step* by composing all the pertinent transformations (i.e. head-motion transform matrices, susceptibility distortion correction when available, and co-registrations to anatomical and output spaces). Gridded (volumetric) resamplings were performed using `antsApplyTransforms` (ANTs), configured with Lanczos interpolation to minimize the smoothing effects of other kernels (Lanczos, 1964). Non-gridded (surface) resamplings were performed using `mri_vol2surf` (FreeSurfer).

### 2.6. Regions of Interest

We extracted blood oxygenation level-dependent (BOLD) signal at specific regions of interest (ROIs) along the auditory pathway: cochlear nucleus (CN), superior olivary complex (SOC), inferior colliculus (IC), medial geniculate nucleus (MGN), Heschl’s gyrus (HG), planum temporale (PT), planum polare (PP), superior temporal gyrus anterior portion (STGa), and superior temporal gyrus posterior portion (STGp). We used the subcortical region definitions from the atlas recently published by Sitek et al. (2019)4. Cortical regions were defined using the Harvard-Oxford parcellation included in FSL 5.0 and accessed through *nilearn* 0.5.2 (Abraham et al., 2014). ROI definitions included both left and right hemispheres. A simple General Linear Model (GLM) sound vs no-sound contrast was calculated using *nistats* 0.0.1b1 to select cortical voxels that respond to sound for subsequent analysis. Nilearn’s `NiftiMasker` was used to extract multi-voxel activity from each of the ROIs. The masks for the cortical regions took the intersection with the subject’s brain mask, as prepared by *fMRIPrep*, and the map of significant (*p* < .05 uncorrected) voxels in the sound vs no-sound contrast. To improve the signal-to-noise-ratio (SNR), the `NiftiMasker` detrended, standardized, and removed confounding variables (as calculated by *fMRIPrep* and described above).

### 2.7. Convolutional Neural Network Activations

The convnets analyzed here are a subset of those analyzed in (Thompson et al., 2019a). All networks were trained to perform context-dependent phone (triphone) classification. Here we look only at the nine freeze-trained networks, which outperformed all other models in Thompson et al. (2019a). These nine networks consisted of three monolingual networks for each of the three languages (English, Dutch and German) and six transfer networks which were first trained on one language and then freeze-trained on another. In all cases, all parameters were updated for 100 epochs and then the networks were freeze-trained for an additional 100 epochs. Freeze training refers to the procedure by which layers are gradually removed from the set of trainable variables over the course of training and in order of depth. Previous work has shown that freeze training can speed up training (Raghu et al., 2017) and facilitate transfer across related tasks (Thompson et al., 2019a). All networks were of identical architecture and consisted of nine convolutional layers followed by three fully connected layers. The layers were as follows, where triplets specify the filter size and number of feature maps in each convolutional layer and the singletons specify how many units in each fully connected layer: (7, 7, 1024), (3, 3, 256), (3, 3, 256), (3, 3, 128), (3, 3, 128), (3, 3, 128), (3, 3, 64), (3, 3, 64), (3, 3, 64), (600), (190), (9000). The input data were 45-dimensional mel-frequency filterbank features calculated at a rate of one frame every 10 ms.

For every network, the activation in response to the original (unquilted) speech stimuli was recorded. For convolutional layers, the average activation within each feature map was recorded. For fully connected layers, the activation at each unit was recorded. Only the activation in response to every second frame of the audio features was saved. Subsequently, the network activations were segmented according to the same phonetic boundaries and were quilted according to the same segment order that was used when generating the experimental stimuli. This produced 180 sequences of network activations for each network, corresponding the 180 speech quilts presented in the scanner.

### 2.8. CKA Similarity Analysis

CKA is a matrix correlation method, similar to representational similarity analysis (RSA) or canonical correlation analysis (CCA). CKA takes two matrices *X* and *Y* as input: in this case, one for the BOLD responses and one for the convnet responses to the same stimuli. CKA can be expressed as a normalized version of the Hilbert-Schmidt Independence Criterion (HSIC) (Cortes et al., 2012).

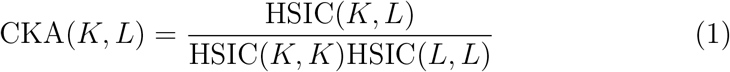

where *K_ij_* = *k*(**x***_i_*, **X***_j_*) and *L_ij_* = *l*(**x***_i_*, **X***_j_*) correspond to two kernels. Gretton et al. (2005) proved that HSIC converges to the population value at a rate of 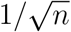. The standard HSIC varies between 0 and 1 where 0 indicates independence between *X* and *Y*. When using a linear kernel, CKA is simply:

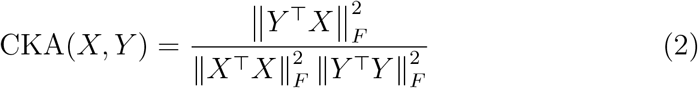

which is equivalent to the RV-coefficient (Robert and Escoufier, 1976).

Here we calculated CKA with a radial basis function (RBF) kernel and an unbiased estimator of the dot product similarity. The choice of the RBF kernel is based on several preliminary network-to-network and brain-to-brain comparisons where the representational hierarchy is known. As described in the Supplemental Material, RBF CKA was most sensitive to the representational similarities of interest. To make CKA less biased, the dot product stimilarity in the standard CKA is replaced with the unbiased HSIC, as described in Song et al. (2007) and as implemented in the Google colab that was released with Kornblith et al. (2019). This unbiased RBF CKA metric varies between −1 and 1.

The matrices *X* and *Y* to be compared must have the same number of rows, corresponding to time points or observations, but can differ in the number of columns, corresponding to voxels or units. Since the temporal rate of fMRI is much slower than that of our acoustic features, temporal rescaling and alignment is required. The preprocessed BOLD timeseries from each ROI and each run were upsampled to match the frame rate of the network activations (one frame every 20 ms) using *pandas* (McKinney, 2010, 2011). This strategy allowed us to preserve the temporal resolution of the network activations without need for summary or binning. The quilted network activations were then aligned to the corresponding BOLD timeseries, setting timepoints when no stimulus was presented to zero. Since the timing of the experimental runs and the stimuli presentation order was different for each subject, this resulted in one matrix per subject per run for each layer of each convnet.

The Glover model of the hemodynamic response function (HRF) (kernel length=32 seconds), as implemented in *nistats* 0.0.1b0, was convolved with the network activations. We extracted and concatenated only the time segments corresponding to the blocks of continuous auditory stimulation from both the fMRI and network activity. The first six seconds of each block were excluded from the analysis to allow for the HRF to ramp up. Thus, the to-be-analyzed fMRI activity does not include the on/off response at the onset of the stimulus blocks. Responses to each block were trimmed to exactly 8599 frames, which, when concatenated, resulted in matrices with 515940 rows for both the fMRI and neural network activity. CKA similarity was then calculated for all ROI-layer pairs

#### 2.8.1. Neural similarity score

To quantify the similarity between a given ROI and network layer, we also calculate the CKA similarity between each ROI and the layers of an untrained network. This untrained network has the same architecture as the trained models, but its parameters have been randomly initialized and never updated. If training has increased the correspondence to the brain, the CKA scores for a trained network should be greater than that of the untrained network. We capture the effect of training on similarity by calculating the difference of standardized CKA scores between a trained network of interest and an untrained network, which we refer to here as the *neural similarity score* for brevity. Within each subject, the CKA scores are standardized using the mean *μ_s_* and standard deviation 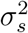. calculated over all models and ROI-layer pairs. The CKA scores of the untrained network are standardized using the same mean and standard deviation. The neural similarity score 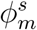 is a difference of z-scores which reflects the similarity achieved by model *m* in subject *s* relative to the untrained model.

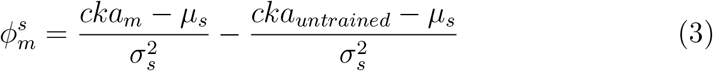

Thus a neural similarity score of 1 indicates that the similarity achieved by the trained model is 1 standard deviation greater than that achieved by the untrained network. As previous work has shown, it is crucial to compare trained networks to a random network to verify that the observed similarity can be attributed to the optimization and is not inherited from the similarity of the input features and/or architecture alone (Kell et al., 2018, Cadena et al., 2019).

## 3. Results

We calculated the CKA similarity for each network, subject, and ROI-layer pair. The results of these analyses can be summarized in similarity matrices whose rows correspond to layers of a network and whose columns correspond to the auditory ROIs. Figure 1 shows the grand mean similarity matrix (left), the mean similarity matrix for the untrained network (middle), and the mean neural similarity score matrix (right). Training increased network similarity to the auditory ROIs, as evidenced by the fact the the neural similarity scores for the trained layers are all positive (Figure 1c). However, we find no evidence of a shared hierarchy, which would manifest itself as a diagonal pattern of high neural similarity scores where shallow layers are more similar to early ROIs and deeper layers are more similar to later ROIs. This hypothesized diagonal pattern also does not occur in the raw CKA similarity scores, neither for the trained nor untrained networks (Figure 1a-b). Instead, for all ROIs, the first fully connected layer (fc1) achieves the highest raw CKA similarity and the highest neural similarity score. This pattern does not occur in the similarity matrix for the untrained network, suggesting that it was introduced by training and not by the architecture.

**Figure 1:**
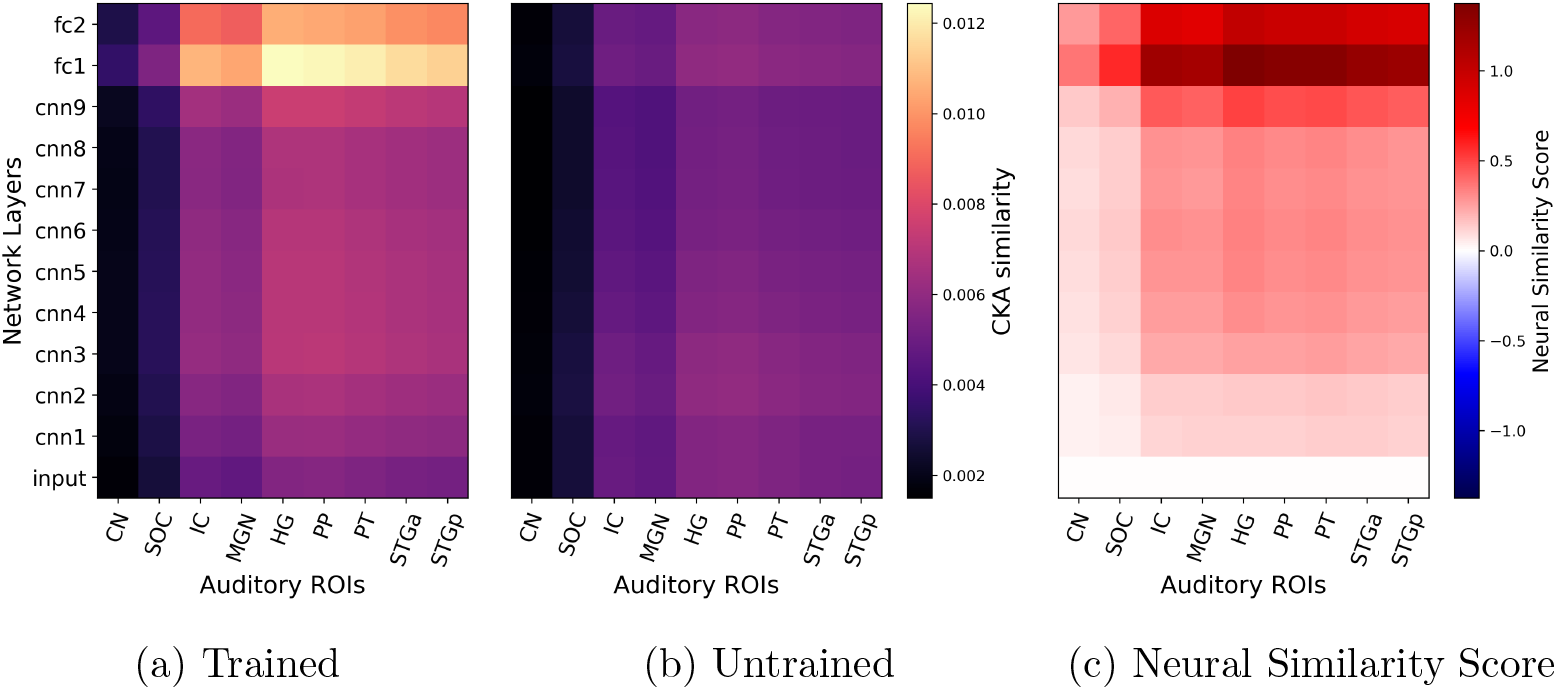
Grand Average Similarity. No shared representational hierarchy is observed. (**Left**) Raw CKA similarity averaged over participants and networks. (**Middle**) Raw CKA similarity for the untrained network, averaged over participants. (**Right**) Neural similarity score averaged over participants and networks. The similarity matrix contains no negative values, showing that training increased correspondence, but there is no diagonal pattern to indicate a shared hierarchy. Instead, for all ROIs, the first fully connected layer (fc1) is most similar.

We calculated the average neural similarity score matrix for each network to investigate how the different training curricula would affect the correspondence. Figure 2 displays nine similarity matrices arranged in a grid. The monolingual models, which were only ever trained on one language, are along the diagonal of the grid. The off-diagonal matrices correspond to the transfer networks which were first trained on one language and subsequently freeze trained on another. The patterns observed in the grand average are largely replicated in the network-specific similarity matrices. Layer fc1 generally achieves high neural similarity scores and none of the networks show any clear evidence for a shared hierarchy. The neural similarity score for layer fc2 is near or below 0 for the monolingual networks but well above zero for the transfer networks. Receiving training on two languages rather than one increased the correspondence between layer fc2 and the auditory ROIs.

**Figure 2:**
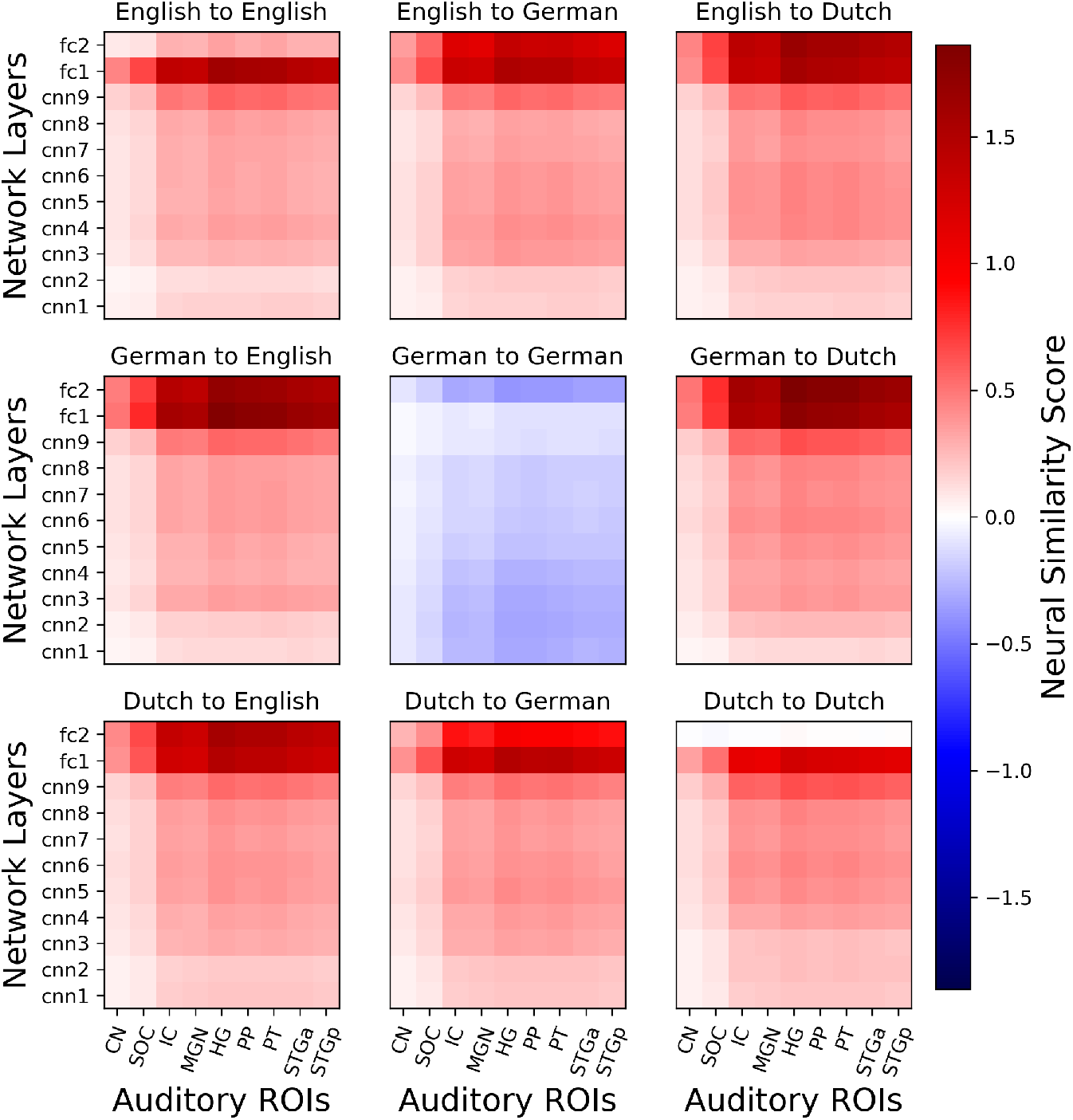
Average Neural Similarity Score. Each similarity matrix shows the effect of training on CKA similarity averaged over the six participants. The subtitles of the form “Language 1 to Language 2” indicate that the network was first trained on Language 1 and then freeze trained on Language 2. Training generally increased the correspondence between brain and networks. Layer fc1 shows the highest neural similarity score and there is little evidence for shared hierarchy (no diagonal pattern). In some layers of certain networks, training did not affect or actually reduced the ROI-layer similarity (shown in white and blue). Layer fc2 yields greater neural similarity for the networks that were trained on two languages, which also performed better on the triphone recognition task.

We hypothesized that the differences between models observed in Figure 2 may be related to the models’ accuracy on the phone classification task on which they were trained. In Figure 3, we plot the peak neural similarity score as a function of triphone classification accuracy. The lines show the linear regression fit for each language-subject pair. All slopes are positive, indicating a positive relationship between model accuracy on the speech recognition task and the peak similarity with the human auditory pathway.

**Figure 3.**
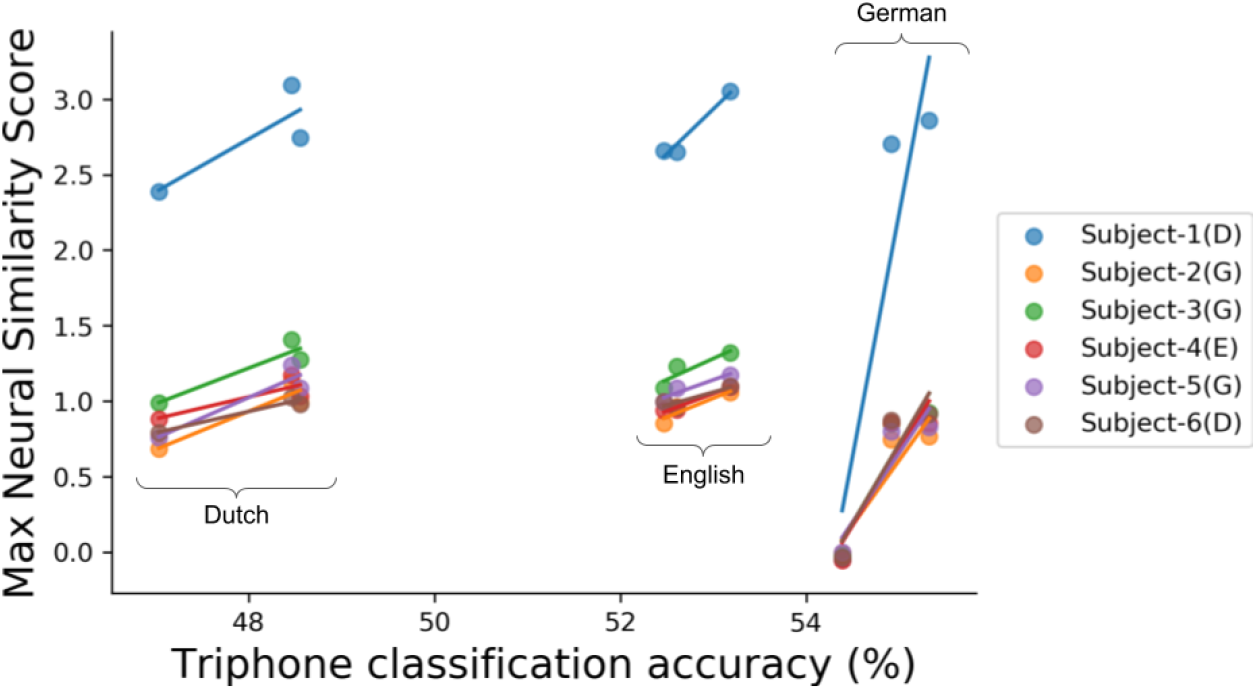
Peak Neural Similarity Score vs Model Accuracy. There are nine points per subject for the nine different network models. Lines show the linear regression fit to the three models (one monolingual and two transfer) for each language and subject. Triphone classification accuracy indicates the top-1 test accuracy achieved by each model. For all language-subject pairs, there is a positive relationship between model accuracy and the correspondence to the human brain. However the effect is largest for the German models, owing to the lesser neural similarity score for the German monolingual model. Parenthetical in the legend indicate the native language of each subject. The regression statistics are reported in the Supplemental Information.

## 4. Discussion

Our experimental results clearly demonstrated that training our convnets on the triphone recognition tasks increased their representational similarity to the collected auditory fMRI activity. This demonstrates that our experimental design and analysis was sufficiently sensitive to reveal training-related effects on representational similarity. However, unlike the previous results of Kell et al. (2018) and Güçlü et al. (2016), this similarity did not manifest in a pattern of shared hierarchy; shallower layers were not most similar to early regions and deeper layers were not more similar to later regions. Instead, the first fully-connected layer, fc1, achieved the highest similarity score across all ROIs, followed by the second fully-connected layer, fc2.

This apparent discrepancy may be best explained by reference to the different cost functions employed and stimuli classes presented. In fact, our results are not inconsistent with previously reports of shared hierarchy. Rather, our work constitutes a stricter test of the shared hierarchy hypothesis and our results suggest the limits of such claims. While we focused specifically on the purely acoustic transformations between spectrogram features and triphones for exclusively speech stimuli, both Kell et al. (2018) and Güçlü et al. (2016) trained networks on tasks at a higher level of abstraction such as word and musical genre recognition and used on a wide variety of natural sounds, effectively analyzing a broader span of auditory features from low-level spectral features up to high-level semantic categories. Recall that the primary evidence of shared representational hierarchy in Kell et al. (2018) was a relatively coarse grain distinction between primary auditory cortex, which was better predicted by shallower layers and secondary auditory cortex, which was better predicted by deeper layers. It is possible that we may have also found a similar distinction had we trained our networks to recognize words. Future work will need to continue to probe the granularity of any shared representational hierarchy, for example by testing the shared hierarchy hypothesis on subsets of network layers.

There is a large diversity of experimental design and analysis approaches employed for the evaluation of representational models. We were inspired by previous fMRI studies which used continuous acquisition during continuous stimulation, for example natural movies, as in the studyforrest dataset. It’s been shown that single trial (i.e. without repetition) measurements during movie watching contain sufficient information to train successful decoding models (Hu et al., 2017) and that functional alignment across subjects based on such single trial measurements can improve decoding performance relative to single-subject decoding (Haxby et al., 2011, Bazeille et al., 2020). Experimental designs of this type sacrifice reliable responses to individual conditions in favor of maximizing the diversity of stimuli presented (which aids generalization) and the number of brain volumes collected. Similarity analyses like CKA benefit from a large number of observations differently than a classical GLM contrast analysis where a robust, reliable response to a small number of conditions is most important. In this way, the optimal design for a similarity analysis may be similar to that of functional alignment. In order to align two representational spaces, either between two brains or between model and brain, the stimulus trajectory should maximally explore the stimulus space of interest. This is why we opted for a continuous stimulation paradigm and approximately two hours of unique speech stimuli, in contrast to previous studies which presented a much smaller number of sounds and analyzed responses averaged over several repetitions. A systematic comparison of different experimental design and analysis methods is needed to tease apart the effect of such choices.

We found that all layers were most similar to fc1 on average. Kell et al. (2018) similarly found that the median variance explained across auditory cortex was maximal at deep but not the deepest layers. This common observation may be related to the notion of dimensionality expansion and compression in DNNs. Recent work describes a two-stage process by which trained DNNs perform a task. The first stage, which might be call ‘feature extraction’, is characterized by increasing intrinsic dimensionality (dimensionality expansion) in the early layers of the network. The second, dimensionality compression, is characterized by decreasing intrinsic dimensionality in the last layers of the network, as the network projects the data to a low-dimensional manifold from which the target can be linearly decoded (Recanatesi et al., 2019, Ansuini et al., 2019). Our layer fc1 may be the last ‘expansion’ layer before the ‘compression’ of the final layers. From Thompson et al. (2019a), we know that layer fc1 is at the barrier between the intermediate layers which are largely transferable between languages, and the final layers which are highly task specific. In Thompson et al. (2019b), layer fc1 was the deepest layer to show a high degree a similarity in networks trained on different languages. The last layers of networks trained on narrowly defined tasks such as triphone recognition may simply learn representations that are more task-specific than any representations employed by the human brain, whose ultimate goal during speech listening is typically natural language understanding, not phoneme recognition. However, fc2 was also found to be relatively similar, but only for the models which were trained on two languages rather than one. These networks benefited from twice the amount of training data as the models trained on only one language and displayed superior generalization as a result. Our analysis revealed that these more generalizable, less language-specific penultimate representations were also more similar to activity in the auditory brain.

Alternative architectures, cost functions, training procedures, or measurement modalities may be required to achieve a layer-to-ROI correspondence for low-level acoustic speech features. Given the low temporal-resolution of fMRI and the temporal nature of sound, incorporating faster measurements such as electroencephalography, magnetoencephalography, or electrocorticograpahy may reveal common patterns that cannot be detected with fMRI. Future work may want to explore non-convolutional model architectures as there are a number of reasons why convnets may not be ideal architectures for audio spectrogram features. Auditory objects display differently in spectrograms than visual objects in images. In particular, auditory objects tend to be less local than visual objects; the part of the spectogram corresponding to a particular sound object is often distributed across several frequencies and time points. Additionally, auditory objects do not occlude each other as visual objects in images do. Instead, overlapping auditory objects in a spectrogram will combine additively. In this way, the inductive bias of convolutional filters is less appropriate for traditional spectrogram-like features (Wyse, 2017) and thus perhaps less likely to yield brain-like representations. Recurrent or autoregressive architectures, which have been very successful in audio synthesis (Oord et al., 2016), may be ideal candidates to investigate in future work.

## Acknowledgments

This work was supported by NWO Vici-Grant 453-12-002 and the Dutch Province of Limburg, an operating grant from the Canadian Institutes of Health Research (MOP 201309), the Erasmus Mundus Student Exchange Network in Auditory Cognitive Neuroscience, a Mitacs-Accelerate internship, and doctoral scholarships from the Fonds de Recherche du Québec – Nature et technologies and Natural Sciences and Engineering Research Council (CREATE). Speech audio was provided by Nuance Communications.

## Footnotes

1 Tags are descriptive text annotations like genre or instrumentation labels.

2 MRI data will be made available on openneuro.org at publication time.

3 Original sound quilting code can be found here: http://mcdermottlab.mit.edu/downloads.html.

4 Due to the small size of CN and SOC and the difficulty of inter-subject alignment of the brainstem, we cannot be completely certain that the activity we extracted truly corresponds to activity in these small brainstem regions. However, the participants in the present study were also participants in the auditory fMRI sessions reported in (Sitek et al., 2019), providing some assurance that these region definitions are reasonable.

