## Supplemental Information for "Training neural networks to recognize speech increased their correspondence to the human auditory pathway but did not yield a shared hierarchy of acoustic features"

### 1. Validating CKA

As mentioned in the main text, we were motivated to use centered kernel alignment (CKA) based on previous work from (Kornblith et al., 2019) showing that CKA was the only metric among several competing methods to accurately identify corresponding layers in two identical networks, differing only in their random initialization. To further validate CKA for our particular application and to select which kernel to use, we performed a number of similar tests where the ground truth hierarchical structure is known. In all of the ridge regression results in the following analyses, to construct a symmetric similarity value, we report the mean variance explained ($R^2$) averaged over two regression fits ($X$ regressed onto $Y$ and $Y$ regressed onto $X$).

#### *1.1. Network to Network Comparison*

The simplest test of whether a similarity metric is sensitive to the representational hierarchy in neural networks is simply to calculate a self-similarity matrix by comparing every layer in a network to every other layer in the same network. This is what's shown in the top row of Figure S1, where the English monolingual model is chosen as a representative example. A metric that is sensitive to the network's representational hierarchy should find neighboring layers similar, i.e. the density of high similarity should be near the diagonal. All metrics tested, CKA with a linear kernel, CKA with an RBF kernel, projection weighted CCA (Morcos et al., 2018), and ridge regression (as implemented in scikit-learn (Pedregosa et al., 2011)), display this pattern to some degree, as evidenced by the fact that the top-left and bottom-right corners are the darkest regions, but RBF CKA displays the expected pattern most smoothly. In particular, comparing linear and RBF CKA, we notice that the similarity between the deepest layers, fc1 and fc2 is better captured by RBF CKA.

We applied the same four metrics to compare the activations of the networks trained on different languages. The English to Dutch comparison is shown in the bottom row of Figure S2 as a representative example. Here we again expect neighboring layers to be similar, but also we want to test whether the metrics find corresponding layers to be most similar. pwCCA fails completely to identify corresponding layers as most similar. Among the other metrics, RBF CKA shows the clearest distinction between corresponding/neighboring and distant layers.

#### *1.2. Brain-to-Brain Comparisons*

Similar to the network-to-network comparisons above, we can also calculate the self-similarity of the BOLD activity by computing all pair-wise comparisons between the ROIs of a particular subject. Given the known connectivity of these regions, we expect the activity in some regions to be

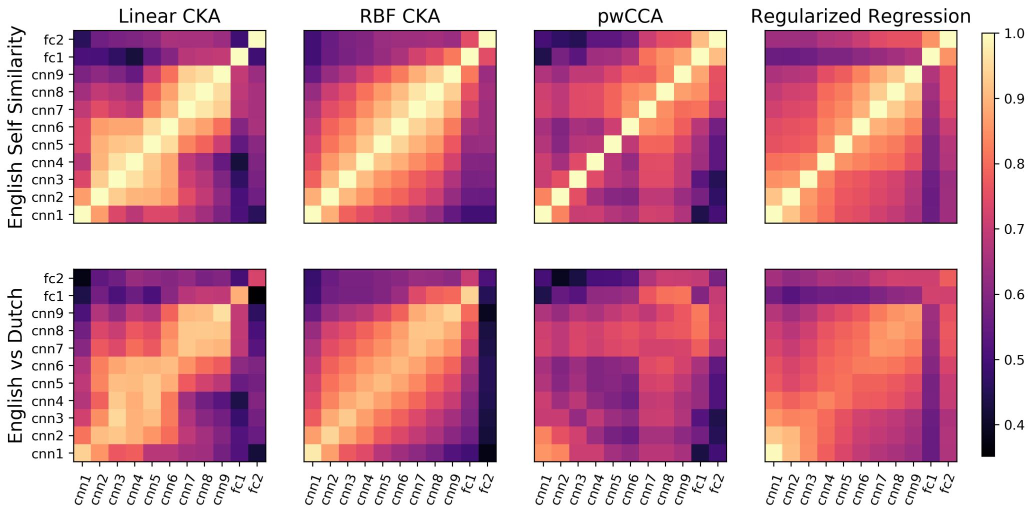


Figure S1: **Network-to-Network Comparisons** CKA with an RBF kernel most clearly displays the similarity of neighboring layers in the English mono-lingual network (top row), and between corresponding layers of two networks trained on different languages (bottom row). Additional metrics: pwCCA - projection weighted mean canonical correlation coefficient; Regularized Regression - variance weighted variance explained, averaged over X predicts Y and Y predicts X.

more similar than others. For example, we expect cortical regions to be more similar to themselves than to subcortical regions, and we would expect HG to be similar to the regions adjacent to it on the superior temporal plane (PP and PT). We also expect PP to be more similar to the anterior portion of the STG, PT more similar to the posterior portion and STGa and STGp similar to each other based on their proximity and connectivity. The top row of Figure S2 shows the mean BOLD self-similarity matrices as calculated with linear CKA, RBF CKA, regularized kernel CCA (with Gaussian kernel)[1] (Bilenko and Gallant, 2016), and ridge regression. To obtain a reasonable computational time for the CCA analysis, we averaged over only the top 50 canonical coefficients. With only these 50 coefficients, regularized kernel CCA captures the expected similarity patterns least well. The other three metrics reveal relatively similar patterns with the one exception that regularized regression shows similarity between cortical and subcortical regions where the CKA methods do not. Therefore, we again conclude that CKA is most sensitive to the known representational hierarchy, but this time find not appreciable difference between the two kernels.

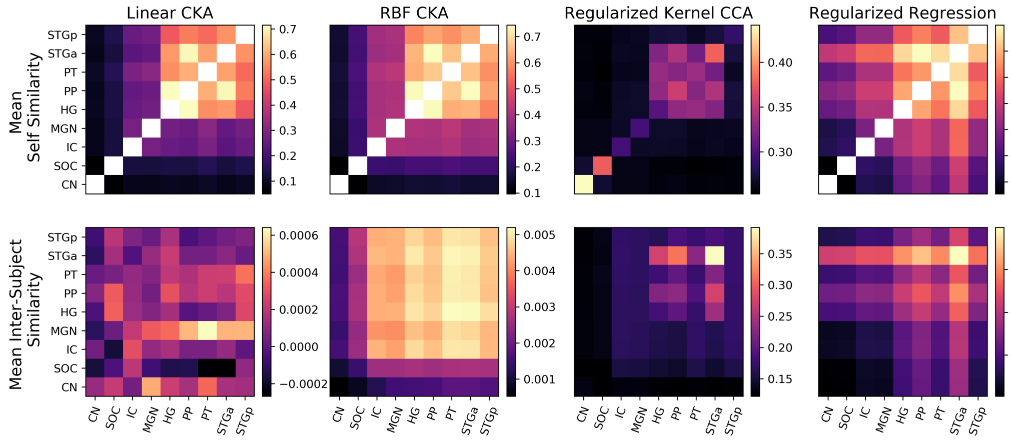


Figure S2: **Brain-to-Brain Comparisons** RBF CKA better reflects expected similarity patterns than linear CKA. White indicates a value of 1. Colour bar adjusted to begin a next lowest value for better visibility. Note: all matrices plotted on their own scales. Other metrics: regularized kernel CCA - variance explained; regularized regression - variance explained averaged over Y predicts X and X predicts Y.

---

[1] Regularized kernel CCA is more appropriate than pwCCA for the small number of BOLD volumes compared to the number of observations that can be made of the neural networks.

Lastly, our most conservative and challenging validation of CKA compares ROIs from different participants. These results should be considered with the caveat that we analyzed single trial BOLD activity, making the comparison across participants fraught with low signal-to-noise ratio. While none of the tested metrics clearly demonstrates corresponding ROIs to be most similar, some metrics fail to reveal this pattern more disastrously than others. Linear CKA resulted in a non-smooth, structureless similarity matrix and very low similarity values. Regularized kernel CCA and ridge regression both identified STGa as being similar across subjects, but yielded similarity values in the same range as for the within-subject comparison, suggesting that these results may be highly overfit. Alternatively, as desired, RBF CKA yields similarity values in a lower range than that of the corresponding self-similarity matrix. RBF CKA also finds cortical regions to be most similar to themselves and is generally smooth, suggesting that the similarity values are not random. Taken together, the network-to-network and brain-to-brain self-similarity and inter-model/inter-subject similarity results further justify our use of RBF CKA in the network-to-brain comparisons of interest.

### 2. Regression Statistics

Table S1 contains the mean slopes for each language and corresponding standard deviation over subjects for the regression described in section 3.

Table S1: **Network Accuracy vs. Peak Neural Similarity Score** Summary of the relationship between network accuracy on the triphone recognition task and peak neural similarity score, as depicted in Figure 3. There was a consistent positive relationship between peak neural similarity score and triphone classication accuracy.

| Language | Mean slope | Standard deviation of slope |
|---|---|---|
| English | 0.29 | 0.14 |
| German | 1.34 | 0.84 |
| Dutch | 0.23 | 0.07 |
